# Regulation of tau’s proline rich region by its N-terminal domain

**DOI:** 10.1101/633420

**Authors:** Kristen McKibben, Elizabeth Rhoades

## Abstract

Tau is an intrinsically disordered, microtubule-associated protein with a role in regulating microtubule dynamics. Despite intensive research, the molecular mechanisms of tau-mediated microtubule polymerization are poorly understood. Here we use single molecule fluorescence to investigate the role of tau’s N-terminal domain (NTD) and proline rich region (PRR) in regulating interactions of tau with soluble tubulin. Both full-length tau isoforms and truncated variants are assayed for their ability to bind soluble tubulin and stimulate microtubule polymerization. We describe a novel role for tau’s PRR as an independent tubulin-binding domain with polymerization capacity. In contrast to the relatively weak tubulin interactions distributed throughout the microtubule binding repeats (MTBR), resulting in heterogeneous tau:tubulin complexes, the PRR binds tubulin tightly and stoichiometrically. Moreover, we demonstrate that interactions between the PRR and MTBR are reduced by the NTD through a conserved conformational ensemble. Based on our data, we propose that tau’s PRR can serve as a core tubulin-binding domain, while the MTBR enhances polymerization capacity by increasing the local tubulin concentration. The NTD negatively regulates tubulin-binding interactions of both of these domains. This study draws attention to the central role of the PRR in tau function, as well as providing mechanistic insight into tau-mediated polymerization of tubulin.

**Significance Statement:** Tau is an intrinsically disordered, microtubule associated protein linked to a number of neurodegenerative disorders. Here we identify tau’s proline rich region as having autonomous tubulin binding and polymerization capacity, which is enhanced by the flanking microtubule binding repeats. Moreover, we demonstrate that tau’s N-terminal domain negatively regulates both binding and polymerization. We propose a novel model for tau-mediated polymerization whereby the proline rich region serves as a core tubulin-binding domain, while the microtubule binding repeats increase the local concentration. Our work draws attention to the importance of the proline rich region and N-terminal domain in tau function, and highlights the proline rich region as a putative target for the development of therapeutics.

## Introduction

Tau belongs to a family of microtubule associated proteins that generally function to modulate microtubule stability and dynamics (1, 2). The deposition of aggregates of tau is linked to a number of neurodegenerative disorders, collectively known as tauopathies [reviewed in (3)]. Cell death is thought to arise both from the process of aggregation as well as from the loss of functional tau and subsequent destabilization of microtubules (3–5).

Tau is an intrinsically disordered protein and it appears to remain largely disordered even upon binding to soluble tubulin (6) or microtubules (7, 8). *In vitro*, tau decreases the critical concentration for tubulin polymerization and regulates microtubule growth rates, catastrophe frequency and recovery (9–11). More recently, tau has been observed to sequester soluble tubulin during liquid-liquid phase separation, leading to microtubule polymerization and bundling (12). It was proposed that this phenomenon may underlie the initiation of microtubules in the axons of neurons.

In the brain, there are six different isoforms of tau, arising from alternative splicing, resulting in the presence of 0, 1 or 2 inserts in the N-terminal domain (NTD) and 3 or 4 repeats within the microtubule binding region (MTBR) (Fig. 1) (13). These isoforms are developmentally regulated with varying distributions of isoforms across developmental stage, cell type and cellular location (3, 13). The microtubule binding region (MTBR) is the best-studied region of tau, both because it contains the 31 or 32 residue long eponymous repeat sequences that are important for binding to microtubules (14, 15), but is also forms the core of aggregates in tauopathies (Fig. 1) (16–18). More recent studies have focused on the role of R’, the ~25 residues C-terminal to the MTBR, a highly conserved sequence sometimes referred to as a pseudo-repeat (Fig. 1) (19–22). N-terminally flanking the MTBR is the proline rich region (PRR) composed of ~25% prolines across two sub-regions, P1 and P2 (Fig. 1). The addition of P2 and R’ to MTBR fragments increases microtubule binding and stimulates polymerization (9–11, 19, 20, 22, 23). The NTD together with P1 is thought to regulate binding to and spacing of microtubules (9, 24–27), and mediate interactions with other cellular partners such as signaling proteins [reviewed in (28)] or the plasma membrane (29).

**Fig. 1.**
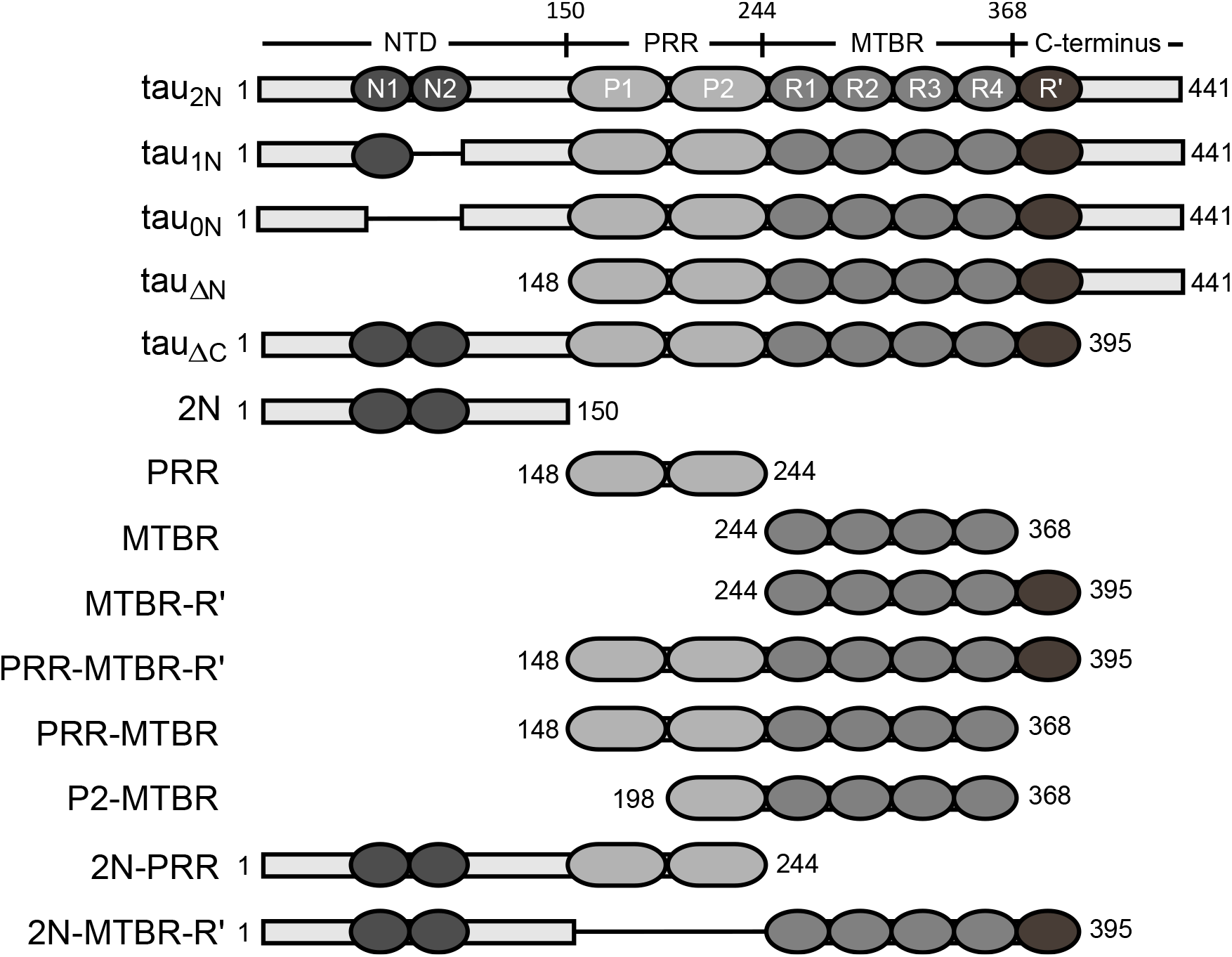
Schematics of tau constructs. The uppermost schematic is the longest tau isoform, tau_2N_. The domains and corresponding residues that delineate them are marked: N-terminal domain (NTD) with N-terminal inserts (N1, N2), proline-rich region (PRR) with sub-regions (P1, P2), microtubule binding repeats (MTBR) with four imperfect repeat sequences (R1-R4) flanked by the pseudo-repeat R’ and the C-terminus. Below are the additional tau isoforms and truncated variants used in this study. All numbering of residues throughout the manuscript is based on tau_2N_.

Despite intense interest in tau, the molecular details of its numerous proposed functions remain relatively obscure. This is in part due to the challenges that arise from its lack of stable structure (30), and that it does not seem to form well-defined stoichiometric complexes with tubulin (20). To illustrate, it was demonstrated more than 20 years ago that P2 (9, 10, 22) greatly enhanced tau binding to microtubules and its ability to polymerize tubulin (31), yet this region of tau is not observed in a recent structure of microtubule-bound tau (7). It may be that the PRR enhances binding indirectly through interactions with the MTBR (22), or that bound PRR is too disordered and dynamic on the microtubule surface to be resolved by EM. These apparently diverging observations, and the need to reconcile them, highlights the requirement for studies of tau function that look beyond the MTBR.

Here, we investigate the role of the NTD and PRR in regulating tau’s interactions with soluble tubulin. Single molecule Förster resonance energy transfer (smFRET) and fluorescence correlation spectroscopy (FCS) of the full-length, N-terminal variant isoforms that contains four MTBR repeats (tau_2N_, tau_1N_, and tau_0N_) were used to monitor binding and probe tau’s conformation in tau:tubulin assembles. We found that in the absence of tubulin the NTD interacts with the PRR and MTBR through a conserved conformational ensemble. The NTD negatively regulates binding to soluble tubulin and subsequently slows polymerization. Strikingly, we find that the isolated PRR is capable of both stoichiometric binding to, and polymerization of, soluble tubulin. The presence of the NTD dramatically reduces the binding and polymerization capacity of the PRR. Based on our results, we propose a model where the PRR serves as a core tubulin binding domain of tau, with both binding and polymerization capacity enhanced by the MTBR and R’, and reduced by the NTD.

## Results

### Conformational ensemble of tau’s NTD/PRR/MTBR is conserved across isoforms

In solution, the N-terminus of tau makes relatively close contacts with both the MTBR and the C-terminus (32), which are lost when tau binds soluble tubulin (6). We used smFRET to assess how the N-terminal inserts impact tau’s solution conformational ensemble. Full-length tau isoforms were labeled with donor and acceptor fluorophores at sites spanning domains of interest (Fig. 2A). The mean energy transfer efficiencies, ET_eff_, were converted to experimental root-mean-square (RMS_exp_) distances using a Gaussian coil model (see *SI Appendix* for details). For constructs probing the C-terminus, tau^291-433^, as well as the PRR, tau^149-244^, all three isoforms gave rise to comparable RMS_exp_ values (Figs. 2B,C and S1B,C and Table S2); this was expected, as the number of residues encompassed by the probes is the same for all three isoforms. The constructs probing the N-terminal domain (NTD), tau^17-149^, also exhibited predicted behavior in that the presence of each N-terminal insert resulted in an increase in the RMS_exp_ (Fig. 2B,C and Table S2). Interestingly, constructs whose labels span the NTD through the PRR, tau^17-244^, or the NTD through part of the MTBR, tau^17-291^ had comparable ET_eff_ histograms, and thus RMS_exp_ values, in solution, despite an increase of up to 60 residues between isoforms (Figs. 2B,C and S1B,C and Table S2). The similar inter-domain distances suggest homologous conformational ensembles between isoforms. Moreover, these ensembles are relatively compact. To illustrate, the RMS_exp_ values of the tau^17-291^ isoforms were of similar magnitude to those of tau^149-244^ despite being 120 to 180 residues longer (Table S2). The RMS_exp_ values for the constructs probing the entirety of the isoforms, tau^17-433^, were also nearly equivalent, consistent with the tau^17-291^ and tau^291-433^ RMS_exp_ values reported here (Figs. 2B,C and S1B,C and Table S2). Upon binding to tubulin, deviations from scaling behavior were diminished, and all constructs yielded RMS_exp_ values that scale with the number of residues in a manner consistent with an extended, random structure (Fig. S1C) (6, 7, 33).

**Fig. 2.**
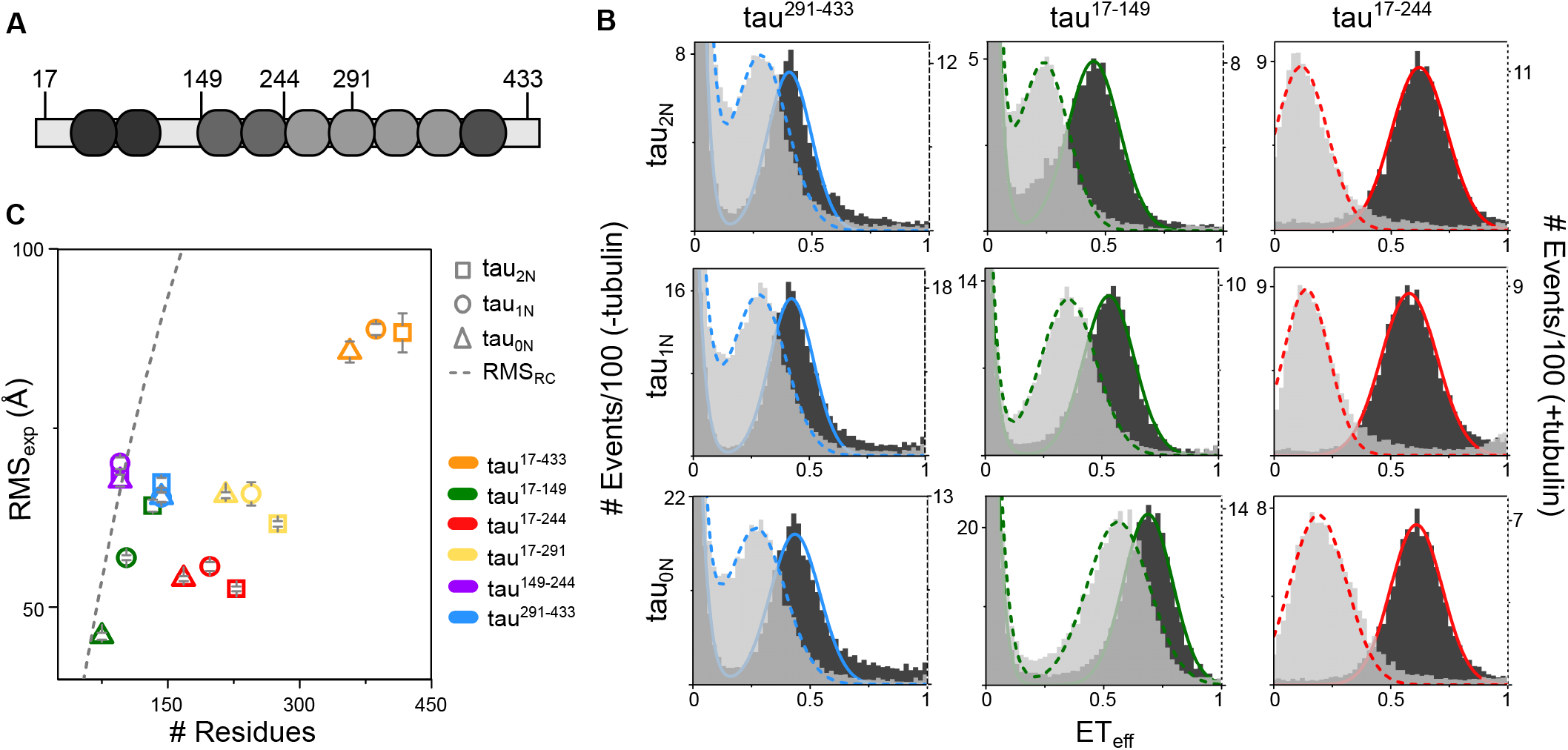
SmFRET of tau N-terminal isoforms. *(A)* Schematic of reference construct tau_2N_ with residues labeled for smFRET measurements are indicated. *(B)* SmFRET histograms of tau_2N_, tau_1N_ and tau_0N_ labeled at residues 291 and 433 (left), 17 and 149 (center), 17 and 244 (right) in the absence (dark: left axis) and presence (light: right axis) of 10 μM tubulin. Histograms for tau labeled at residues 17 and 433, 17 and 291, 149 and 244 are in Fig. S1. Fits to the data as a sum of Gaussian distributions as described in the *SI Appendix* are shown. *(C)* For measurements in the absence of tubulin, the root-mean-square distances (RMS) between labeling positions calculated from a Gaussian chain model (RMS_exp_) are plotted (47). Data are presented as mean ± SD, n≥3 independent measurements. For reference, the RMS calculated for a random coil (RMS_RC_) as in Reference (48) is indicated by the gray dashed line. See Table S2 for numerical values of ET_eff_ mean ± SD, RMS_exp_ ± SD and RMS_RC_ for each construct.

### Tau’s NTD negatively regulates tubulin binding

The conservation of the conformational ensembles across N-terminal isoforms suggests a functional origin. This led us to examine the impact of the N-terminal inserts on tau binding to soluble tubulin. Tubulin binding of fluorescently labeled tau in the presence of increasing concentrations of tubulin was assessed by FCS under non-polymerizing conditions. Both the longest full-length isoform, tau_2N_, and an NTD deletion fragment, tau_ΔN_ (amino acids 149 to 441), bound tubulin as seen by an increase in the normalized diffusion time, τ_norm_, with increasing tubulin concentration (Fig. 3A). However, there are significant differences in the binding curves; tau_2N_ reached its maximum τ_norm_ at ~2.5 μM tubulin, while the τ_norm_ for tau_ΔN_ continued to increase. At 10 μM tubulin, the τ_norm_ of tau_ΔN_ was more than 2x larger than that of tau_2N_ (Fig. 3A). This effect was specific to the NTD. Binding by a C-terminal deletion construct, tau_ΔC_ (amino acids 1 to 395), resembled that of tau_2N_ (Fig. 3A) while a combined N-terminal and C-terminal deletion construct, PRR-MTBR-R’ (amino acids 149 to 395) behaved like tau_ΔN_, (Fig. 3A). These measurements suggest that the NTD of tau reduces or negatively regulates its binding to soluble tubulin, while the C-terminus does not have a significant role. Finally, we measured the tubulin binding by all three N-terminal isoform variants, tau_2N_, tau_1N_ and tau_0N_, and found their binding curves to be comparable (Fig. S2A), indicating that regulation of binding is intrinsic to the NTD and not strongly dependent on the presence or absence of a specific insert.

**Fig. 3.**
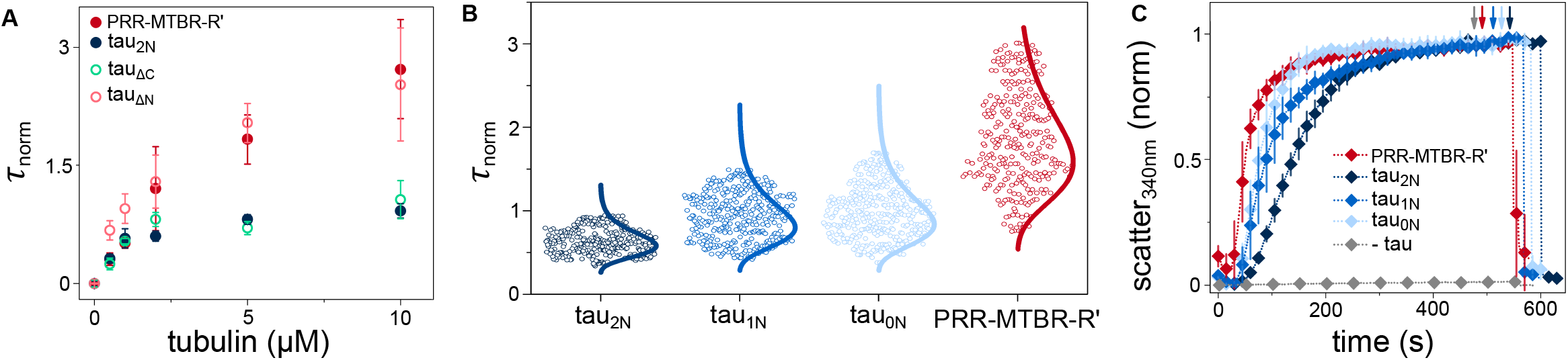
Inhibition of binding and polymerization by the NTD. *(A)* The increase in the normalized diffusion time, τ_norm_, as a function of tubulin concentration reflects binding of fluorescently labeled tau to unlabeled tubulin. Data are presented as mean ± SD, n≥3 independent measurements. See *SI Appendix* for details of data analysis. See Table S5 for numerical values for τ_D_ and τ_norm_ at 10 μM tubulin. *(B)* The autocorrelation curves for tau_2N_, tau_1N_, tau_0N_ and PRR-MTBR-R’ in the presence of 10 μM tubulin were individually fit to obtain a distribution of τ_norm_ values. When the NTD is absent, larger tau-tubulin complexes form as seen by the larger values of τ_norm_. Overlays are lognormal distributions. See Table S3 for descriptive statistics of distributions. *(C)* Tubulin polymerization as measured by scattered light at 340 nm as a function of time. See Table S4 for fit parameters. Data are presented as mean ± SD following normalization, n=3 independent measurements. Arrows indicate depolymerization at 4 °C.

In our prior work, we demonstrated tau forms fuzzy complexes with soluble tubulin consisting of multiple, weakly-associated tubulin dimers (20). Using a similar approach as described in that work, we analyzed the individual autocorrelation curves taken in the presence of 10 μM tubulin in order to assess the heterogeneity of the tau-tubulin complexes (Fig. 3B; details of analysis are in the *SI Appendix*). From this analysis, it was apparent that not only were tau:tubulin complexes formed by PRR-MTBR-R’ on average larger (median diffusion time, τ_median_=2.02 ms) than those formed by any of the full-length constructs (τ_median_=1.29, 1.50, and 1.55 ms for tau_2N_, tau_1N_ and tau_0N_, respectively), but that PRR-MTBR-R’:tubulin complexes also had the largest spread in diffusion times (1.26 to 2.89 ms). These complexes persisted at 300 mM KCl, indicating they were not the result of low salt buffer (Table S5). Analysis of the average brightness of the diffusing species demonstrated that while the full-length constructs typically consisted of a single tau molecule, the PRR-MTBR-R’:tubulin complexes, especially the larger ones, may have included several tau molecules (Fig. S2B,C and Table S6). Together, analysis of diffusion time and brightness of the complexes suggest that the NTD limits both: (1) the average number of tubulin dimers bound to a single tau molecule; and (2) the average number of tau bound to a single tubulin dimer.

The prior work from our lab also demonstrated a positive correlation between the rate of tau-mediated tubulin polymerization and the median diffusion time, and thus the number of tubulin dimers associated, of tau:tubulin complexes (20). That work, however, exclusively used fragments of tau, none of which included the entire PRR or the NTD. Nevertheless, we found that this observation broadly holds for the full-length and fragments studied here: PRR-MTBR-R’ had the fastest polymerization half-time (*t_1/2_*=52 ± 7 s) while the full-length isoforms were all slower (Fig. 3C). Interestingly, tau_2N_ was the slowest (*t_1/2_*=137 ± 9 s), lagging behind tau_1N_ and tau_0N_ (*t_1/2_*=88 ± 13 s and 76 ± 10 s, respectively) (Fig. 3C and Table S4). This, along with the small decrease in the size and heterogeneity of the tubulin complexes formed by tau_2N_ (Fig. 3A), suggests that the N2 insert may have an additional regulatory role in binding and polymerizing tubulin.

### The PRR independently binds and polymerizes tubulin

The reduced binding of NTD containing constructs (Fig. 3) coupled with the conserved conformational ensembles in the NTD/PRR/MTBR constructs observed in the smFRET measurements (Figs. 2 and S1), led us to hypothesize that the NTD may regulate tubulin binding though interactions with the PRR or MTBR. To investigate this hypothesis, we created constructs corresponding to these domains and measured binding by FCS as well as tau-mediated polymerization. Although the MTBR (amino acids 244 to 372) associates with microtubules in the context of the full-length protein or in constructs containing P2 (7, 15), the isolated domain bound only weakly to soluble tubulin (Fig. 4A) and was not capable of polymerizing tubulin (Fig. 4B). The addition of R’, MTBR-R’ (amino acids 244-395), enhanced binding (Fig. 4A) but still did not yield a construct that promoted efficient polymerization (Fig. 4B). Although early studies demonstrated that the MTBR-R’ (9) or even peptides corresponding to the individual MTBR repeats (34) had weak polymerization capacity, 5 to 10-fold more tau was required than the 10 μM used here.

**Fig. 4.**
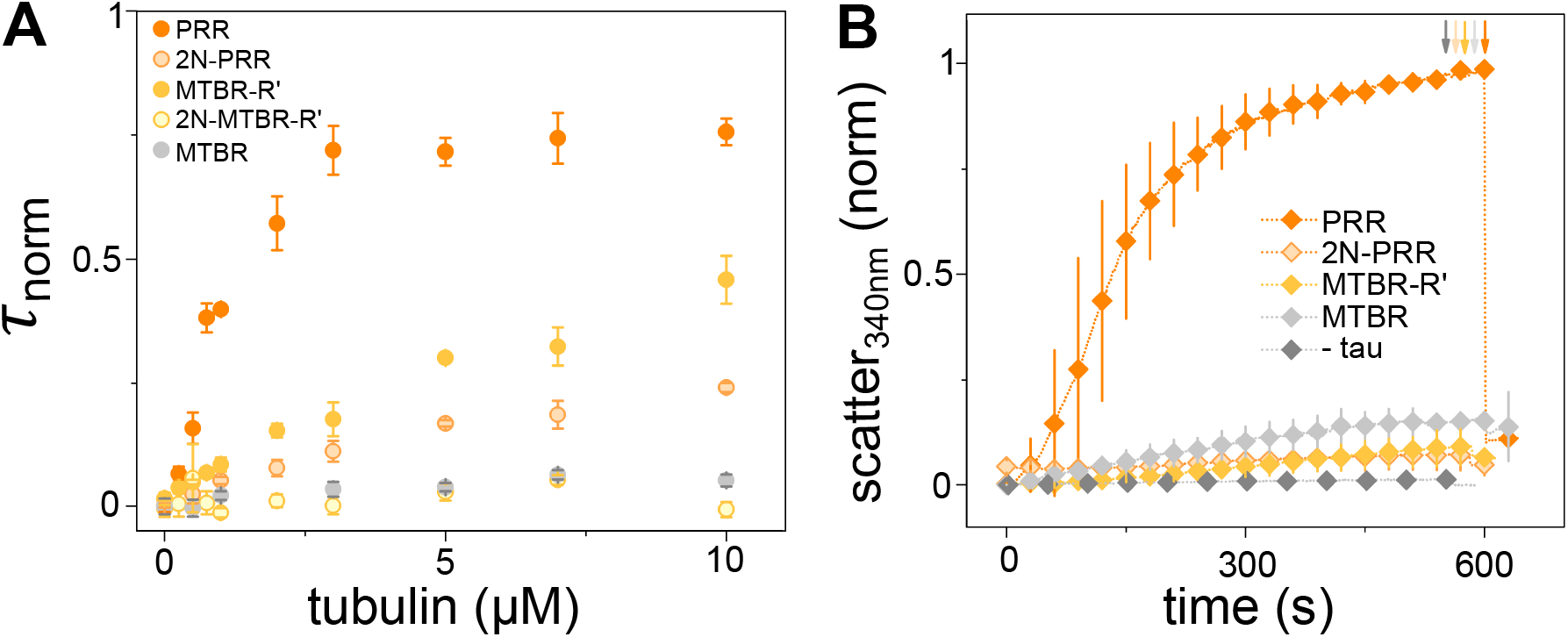
Independent polymerization capacity of the PRR, regulated by the NTD. *(A)* Binding of tau constructs to tubulin as measured by an increase in τ_norm_ as a function of tubulin concentration. Data are presented as mean ± SD, n≥3 independent measurements. *(B)* Tubulin polymerization as measured by scattered light at 340 nm as a function of time. See Table S4 for fit parameters. Data are presented as mean ± SD following normalization, n=3 independent measurements. Arrows indicate depolymerization at 4 °C.

Strikingly – and surprisingly – the isolated PRR (amino acids 149 to 244), bound tubulin tightly when compared to the MTBR and MTBR-R’ measured under the same conditions (Fig. 4A). It also stimulated rapid polymerization of tubulin (Fig. 4B). Fitting the PRR:tubulin binding curve to the Hill equation yields n=1.7 ± 0.2, reflecting cooperativity, and with an apparent K_D_ ≈ 900 nM (Fig. S3). Furthermore, unlike constructs where the PRR is coupled with the MTBR and/or R’, such as PRR-MTBR or PRR-MTBR-R’, the PRR demonstrated saturable binding and did not form large tau:tubulin complexes (Fig. S4). Based on saturation diffusion time measured for PRR:tubulin at 10 μM tubulin, we estimate there to be ~2 binding sites for tubulin dimers in the PRR (see *SI Appendix*). This apparent specificity suggests that formation of tau:tubulin fuzzy complexes arises primarily from the collective binding properties of the PRR and MTBR-R’.

While the PRR and R’ have been identified as enhancing binding and accelerating polymerization *in vitro* (9–11, 19, 20, 22, 23, 35, 36) these features have not previously been observed for the isolated PRR. This may in large part be due to the widespread use of the K16 fragment consisting of P2 and the 4R MTBR (amino acids 198 to 372, P2-MTBR) (9). The P2-MTBR construct binds to tubulin, however, it does not bind as many tubulin dimers at high tubulin concentrations as PRR-MTBR (Fig. S5). Thus, while the isolated P1 does not bind tubulin strongly (Fig. S6), it does enhance binding and contribute to tau function. Tight binding to tubulin required the presence of both proline rich regions; fragments corresponding to P1 (amino acids 149 to 198) or P2 (amino acids 199 to 244) bound tubulin only weakly (Fig. S6).

### NTD negatively regulates the polymerization capacity of the PRR

Our observation that the PRR bound to and polymerized tubulin independently of the MTBR (Fig. 4), combined with the slower polymerization rate of tau constructs including the NTD (tau_2N_, tau_1N_ and tau_0N_) relative to PRR-MTBR-R’, motivated us to determine the impact of the NTD on interactions of the PRR with tubulin. Tau_2N_ was truncated after the PRR at amino acid 244 (2N-PRR), and binding to soluble tubulin and polymerization were measured. The presence of the NTD dramatically reduced binding (Fig. 4A) as well as significantly diminished tubulin polymerization capability (Fig. 4B). Truncated constructs based on tau_1N_ and tau_0N_, 1N-PRR and 0N-PRR respectively, showed similar binding behavior (Fig. S7). Collectively, these results led us to propose that the binding and – by extension – polymerization capacity of tau is regulated by interactions between the NTD and the PRR, evident by the conserved ensembles observed with smFRET for this domain (tau^17-244^ in Fig. 2). Because the conserved ensembles extend into the MTBR (tau^17-291^ in Figs. 2 and S1), we tested this idea by making a construct lacking the PRR (2N-MTBR-R’ amino acids 1 to 148 fused to 245 to 395). This construct also did not demonstrate appreciable binding to tubulin (Fig. 4A), while that same construct lacking the NTD (MTBR-R’) clearly did (Fig. 4A). As a whole, these results strongly support a functional, regulatory role for the compact, albeit disordered, NTD/PRR/MTBR ensembles observed by smFRET.

## Discussion

Since it was first isolated over 40 years ago (1), both functional and structural studies of tau have primarily focused on the MTBR (7–9, 14, 37). Our current study examines two domains of tau that have been the subject of significantly less scrutiny: the NTD and the PRR. Our discovery that the isolated PRR has the capacity to bind tubulin and polymerize microtubules *in vitro*, and that this function is negatively regulated by the NTD, draws attention to the importance of these two domains in understanding tau function.

The NTD has previously been shown to regulate *in vitro* polymerization of tubulin (9). It is implicated in isoform-dependent spacing of microtubules (24, 25), and removal of the NTD both increases the affinity for microtubules as well as results in the presence of large microtubule bundles (9). Here we observed a similar inhibitory effect in binding to soluble tubulin (Fig. 3A). This inhibition seems to be an effect of the NTD as a whole, rather than resulting from the absence or presence of a specific insert, as only small differences in binding are observed for the 0N, 1N and 2N isoforms in comparison to when the NTD is absent (Fig. 3B). Insight into why the inserts do not have a significant effect in regulating binding is gained from our smFRET measurements which show that the relative dimensions corresponding to the NTD-PRR (tau^17-244^) or NTD-MTBR (tau^17-291^) are independent of the number of inserts (Fig. 2 and S1). This suggests that conserved long-range interactions and/or conformational features of the NTD are important for regulating interactions with tubulin, more so than the inserts themselves. Given that the NTD also significantly reduces the size and heterogeneity of ‘fuzzy’ tau-tubulin complexes (Fig. 3) (20), it follows that the NTD may dynamically shield the weak tubulin binding sites distributed throughout the MTBR and R’ (Fig. 5). Our results suggest this screening is a general function of the NTD that serves as an initial regulatory gate to tau-mediated polymerization, and is largely independent of the individual N-terminal inserts. These results also indicate that the regulatory mechanism of the NTD for tau:tubulin is distinct from that of tau:microtubules, where the relative electrostatics and sterics of different N-terminal isoforms impact the tau:tau interactions required for spacing and bundling of microtubules (9, 24–27).

**Fig. 5.**
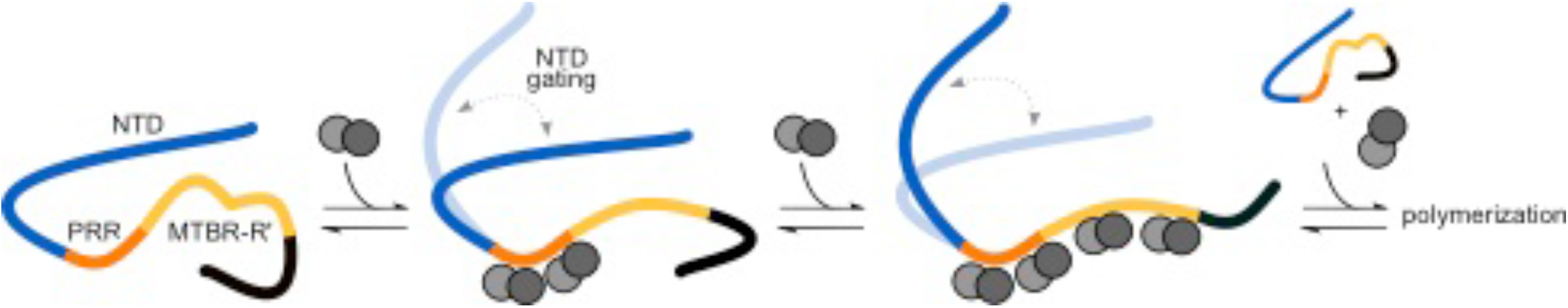
Model for regulation of tau:tubulin interactions. The PRR (orange) binds tubulin tightly and stoichiometrically, negatively regulated by the NTD (blue). The MTBR-R’ (yellow) increases the local tubulin concentration through distributed, weak interactions, enhancing the polymerization capacity of tau. The C-terminus is colored black. Increasing both tau and tubulin concentrations favor polymerization.

Our observation of binding to and assembly of tubulin by the isolated PRR was unexpected, as to our knowledge, there are no prior reports of this in the literature. NMR chemical shifts suggesting binding were measured in the PRR of longer tau fragments in the presence of engineering tubulin constructs (8, 11). For a tau fragment covering part of the PRR, residues 166-244, 1:2 tau:tubulin dimer stoichiometry was observed with taxol stabilized microtubules, but polymerization was not tested (11). We also find tight, saturable stoichiometric binding of 1:2 tau:tubulin dimers. Notably, PRR residues were not observed in the recent cryo-EM structure of microtubule-bound tau (7). It may be that the PRR binds to a region unresolved within the structure, such as intrinsically disordered loops or the tubulin tails, or to a site partially occluded in the microtubule lattice.

The stoichiometric binding to soluble tubulin of the PRR provides a striking contrast with the dynamic, heterogeneous ‘fuzzy’ tau-tubulin complexes formed when the MTBR and R’ were present in the constructs. In particular, tight and specific binding of the PRR may offer an attractive target for drug development relative to the comparatively weak binding by the MTBR-R’ (38). Interestingly, both P2 and R’ were identified relatively early as sequences important for productive tau-mediated polymerization (39). The ‘jaws’ model proposed that targeting of tau to the MT lattice is through these regions, while the MTBR played a catalytic role in assembly (10, 19, 39). However, in our study, the PRR is the only isolated domain which demonstrates any significant tubulin polymerization capacity; this is not seen for MTBR nor MTBR-R’ (Fig. 4). This leads us to propose a variation to that model (Fig. 5). In our model, the PRR serves as the core tubulin binding domain, binding to two tubulin dimers in a critical step towards initiating polymerization. Multiple weak tubulin binding sites in the MTBR and R’ allow for increasing the local concentration of tubulin, resulting in accelerated microtubule growth. The ubiquitous screening by the NTD of both the PRR and the MTBR serves as an initial gating that controls the size of these tau:tubulin ensembles and, consequently, tubulin assembly.

Tau’s interactions with microtubules are regulated by phosphorylation [reviewed in (40) and (41)] and the majority of tau’s 20+ known phosphorylation sites are located in the PRR, including those associated with Alzheimer’s disease (42, 43). Given this, perhaps the relative importance of the PRR in both binding to and polymerizing tubulin should not be so surprising. In contrast, the MTBR is primarily modified by lysine acetylation. Individual or combinatorial effects of these modifications may alter both the binding affinity and the stoichiometry of tubulin binding. There is a least one example of coordinated modifications to tau in the literature: acetylation at residues 280/281 within the MTBR influences phosphorylation at residues S202/205 within the PRR (44). Either of these modifications may in turn influence the interaction of the PRR or MTBR with the NTD, suggesting that regulation of binding may be more complex than simply reducing affinity and stoichiometry, but instead a intricate interplay between the NTD, PRR and MTBR domains.

## Material and Methods

### Protein expression and labeling

Tubulin was purified from fresh bovine brains as described previously (45). Tau constructs were cloned using site-directed mutagenesis or deletion cloning techniques, and purified as described previously (46). Site-specific labeling for FCS and FRET measurements were though introduced cysteines reacted with Alexa Fluor 448 or Alexa Fluor 594 maleimide. See *SI Appendix* for details.

### smFRET and FCS measurements

SmFRET and FCS measurements were carried out on a home-built instrument based on an inverted Olympus 1X-71 microscope (20, 46) or a commercial MicroTime 200 time-resolved confocal microscope (Picoquant). For smFRET, 20-40 pM tau was used; for FCS, 15-25 nM tau was used. Measurements were made in the absence or presence of tubulin at 20 °C under non-assembly conditions in phosphate buffer (20 mM potassium phosphate pH 7.4, 20 mM KCl, 1 mM MgCl_2_, 0.5 mM EGTA, 1 mM DTT). For smFRET, the ET_eff_ was calculated from photon burst intensities and converted to RMS_exp_ distances using a Gaussian polymer chain model (47). For FCS, multiple autocorrelation curves were collected and fit using a single-component 3D diffusion model (20). For heterogeneity analysis, curves were assessed for goodness-of-fit and then filtered until a stable population was reached using iterative filtering of diffusion times. See *SI Appendix* for details.

### Tubulin polymerization

Clarified tubulin was incubated with tau at varying ratios at 4 °C prior to the addition of 1 mM GTP and immediate transfer to phosphate buffer in a cuvette equilibrated to 37 °C. Polymerization was monitored by an increase in the scattered light signal. Normalized curves were fit to a sigmoidal growth equation, Eq. S3. See *SI Appendix* for details.

## Author contributions

K.M. and E.R. designed research. K.M. performed research and analyzed data; K.M. and E.R. wrote the paper.

## Acknowledgements

We would like to thank S. Wickramasinghe, H.Y.J Fung, I. Yannatos and H. Merens for assistance during the various tubulin purifications, and D. Chenoweth for use of his fluorimeter. This work was supported by NIH Institutional Training Grant T32 GM071399 (to K.M.) and NIH Grant AG053951 (to E.R.).

## SI Appendix

### Tubulin purification and handling

Tubulin was purified from fresh bovine brains as described in (1). Purified tubulin was snap-frozen in BRB80 (80mM PIPES pH 6.8, 1mM MgCl_2_, 1mM EGTA). Prior to use, frozen aliquots were rapidly thawed and then clarified at 100,000xg for 6 minutes. BioSpin 6 columns (BioRad) were used to buffer exchange tubulin into phosphate buffer. The tubulin absorbance at 280 nm was converted to concentration using a molar extinction coefficient of 115,000 M^-1^cm^-1^. Tubulin was used within 2 hours following clarification.

### Tau cloning, purification, and labeling

The parent tau plasmid encodes for longest tau isoform, tau_2N_. It includes an N-terminal His-tag with a tobacco etch virus (TEV) protease cleavage site for purification (2). The native cysteines, C291 and C322, are mutated to serine to allow for the introduction of cysteines at desired location for site-specific labeling. Tau_1N_ and tau_0N_ were generated using deletion cloning from the tau_2N_ plasmid. The nicked DNA fragments were fused using T4 DNA ligase (New England Biolabs) and T4 Polynucleotide kinase (New England Biolabs). The remaining tau fragments were generated using either site-directed mutagenesis to introduce stop codons and cysteines, deletion cloning of the remaining tau amino acids within the parent tau vector or a combination of the two techniques.

For all constructs longer than 200 residues, tau protein expression was induced with 1mM IPTG at OD ~0.6 overnight at 16 °C. For constructs <200 residues, tau protein expression was induced with 1mM IPTG at OD ~0.8 for 4-5 hours at 37°C. Purification was based on previously reported methods (2). Briefly, cells were lysed by sonication, and the cell debris pelleted by centrifugation. The supernatant was incubated with Ni-NTA resin (Qiagen or BioRad) and the recombination protein was bump eluted with 500 mM imidazole. The His-tag was removed by incubation lab purified TEV proteinase for either 4 hours at 20 °C (constructs <200 residues) or overnight at 4 °C (constructs >200 residues). Uncleaved protein was removed by a second pass over the Ni-NTA column. Remaining contaminants were removed using size exclusion chromatography on a HiLoad 16/600 Superdex 200 Column (GE LifeSciences). Proteins that did not require fluorescent labeling were buffer exchanged using Amicon concentrators (Millipore) into the final assay buffer of interest, aliquoted and snap frozen for storage at −80 °C. Due to the small size and lack of aromatic residues, P1 and P2 were TEV-cleaved as described above but after the fluorescent labeling (below). All other proteins were labeled following elution from the size exclusion column.

Site specific labeling of tau for FRET or FCS measurements was carried out as described previously (2). Briefly, tau was incubated with 1 mM DTT for 30 minutes, and then buffer exchanged into labeling buffer (20 mM Tris pH 7.4, 50 mM NaCl, and 6 M guanidine HCl). For FRET labeled constructs, the donor fluorophore, Alexa Fluor 488 maleimide (Invitrogen), was added at sub-stoichiometric ratios (0.3-0.5x), and incubated at room temperature for 15 minutes. A 3-fold molar excess of the acceptor fluorophore, Alexa Fluor 594 maleimide (Invitrogen) was added and the reaction was incubated for another 10 minutes at room temperature, and then moved to 4 °C for overnight incubation. For FCS labeled constructs, Alexa Fluor 488 maleimide was added in 3-fold molar excess and incubated at room temperature for 10 minutes, followed by overnight incubation at 4 °C. Labeling reactions were protected from ambient light and with constant stirring; the dye was added dropwise. The labeled protein was buffer exchanged into 20 mM Tris pH 7.4 and 50 mM NaCl and unreacted dye was removed using HiTrap Desalting Columns (GE Life Sciences). Labeled protein was aliquoted and snap frozen for storage at −80°C.

### FCS instrument and data analysis

All FCS measurements were preformed on our home built instrument, as described previously (3). Prior to entering the inverted Olympus 1X-71 Microscope (Olympus), the laser power was adjusted to ~ 5 μW (488 nm diode-pumped solid-state laser, Spectra-Physics) and focused into the sample via a 60x water objective (Olympus). Fluorescence emission from the sample was collected through the objective, separated from excitation light by a Z488RDC long pass dichroic and a 500 nm long pass filter (Chroma). The filtered emission was focused the aperture of a 50 μm diameter optical fiber (OzOptics) coupled to an avalanche photodiode (Perkin-Elmer). A digital correlator (FLEX03LQ-12, Correlator.com) generated the autocorrelation curves.

Measurements were made in 8-chamber Nunc coverslips (Thermo-Fisher) passivated by incubation with (ethylene glycol)poly(L-lysine) (PEG-PLL)(4). The labeled tau (15-25 nM) and tubulin (concentrations vary) were incubated in chambers for 5 minutes prior to measurement. Unless otherwise noted, all FCS experiments were carried out in phosphate buffer (20 mM phosphate pH 7.4, 20 mM KCl, 1 mM MgCl_2_, 0.5 mM EGTA, 1 mM DTT) at 20 °C. Multiple (20-40) 10 second autocorrelation curves were collected per sample and fit to a single-component 3D diffusion equation (Eq. S1):

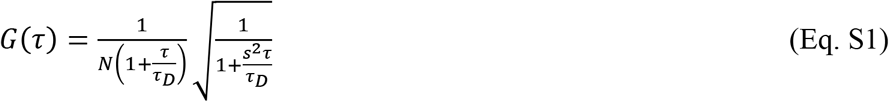

where *G(τ)* is the autocorrelation function as a function of time (*τ*), *τ_D_* is the translational diffusion time of the labeled molecules and *N* is the average number of fluorescent species. For our instrument, the ratio of the radial to axial dimensions of the focal volume (s) was determined to be 0.2 and consequently fixed for analysis. The recorded intensity trace is divided by N to give counts per molecule (CPM) in kHz.

For some tau constructs, high tubulin concentrations (>2 μM) result in the formation of large, bright species (Figs. 3, S2 and S4). These species are not present in the traces of protein in the absence of tubulin (Figs. S8 and S9A). A prior study from our lab demonstrated these species are tau:tubulin specific, electrostatically sensitive and reversible (3). For P2-MTBR, increasing the KCl concentration in our phosphate buffer to 300 mM – previously seen by NMR to disrupt interactions between the PRR and tubulin – results in disassembly of the larger species (Table S5) (5). In the case of PRR-MTBR-R’, these species persist even at 300 mM KCl suggesting the binding is either tighter or has a more hydrophobic character (Table S5).

The individual autocorrelation curves arising from these larger assemblies disproportionally weight the averaged autocorrelation curves used in the analysis described above (Table S3 and Fig. S9). Working under the premise that removal of these outliers would allow for a more meaningful analysis of the majority tau:tubulin complexes, we developed an algorithm to remove aberrant curves, broadly following the approach we described previously (3). Individual autocorrelation curves were fit with Eq. S1 and assessed the goodness of fit using least-squares *X^2^* = [G(τ)_fit_ – G(τ)_raw_]^2^ with a tolerance of *X^2^* = 0.0001 for a consecutive run of 75 ms. In other words, if the fit deviated beyond the *X^2^* for more than 75 ms, the curve was discarded. This process removes 99.5% of curves that cannot be accurately fit using Eq. S1. The frequency of these aberrant curves is ~3%.

Autocorrelation curves arising from larger assemblies that pass this initial criterion still skew the data towards slower diffusion times (Table S3). Descriptive statistics of these diffusion times are reported in Table S3 as “pre-filtering”. In some cases, such as PRR-MTBR-R’ the measured τ_D_ could be as large as ~14 ms and up to 4x brighter than unbound tau (Fig. S9B,C). Although of potential interest in another context, these species do not represent the majority of the tau:tubulin complexes of interest here. These outliers were removed in an iterative fashion by testing the individual curves using an Anderson-Darling statistical test for either a lognormal or normal distribution. Diffusion times above or below the interquartile range were removed until a stable population was reached and no more curves were removed from the dataset. We did not: (1) enforce a lognormal or normal distribution on the data-set prior to outlier removal; nor (2) continue or use outlier removal if the population is normal or lognormal after testing. This iterative function is demonstrated for tau_2N_ in the absence (Fig. S9D) or presence of 10 μM tubulin (Fig. S9E). The initial iteration simply tests for normality (seen by the straight line in Fig. S9D for a single iteration). This results the removal approximately ~15-20% of curves that passed the initial goodness-of-fit filtering from the data set (Table S3).

The fit parameters from the individual filtered curves are presented in Figs. 3B (τ_norm_) and S2B (CPM), and the descriptive statistics of these values are listed in Table S3. There is a general correlation showing that tau:tubulin complexes with larger τ_D_s also had larger CPMs, reflecting the presence of multiple tau molecules in these assemblies (Fig. S2C). In order to allow for straightforward comparison between tau isoforms, we also averaged the filtered curves from each independent measurement and fit the average curve with Eq. S1. These τ_D_ values obtained from these fits are reported in Table S5 for saturating points, and are graphed in figures with FCS binding curves (Figs. 3, 4, S2, S3, S5, S6 and S7).

Tau constructs of different lengths have different diffusion times. To allow for straightforward comparison of the extent of binding between the various constructs, the diffusion times for each construct in the presence of tubulin 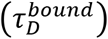 were normalized to that of the construct in the absence of tubulin 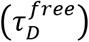 as follows:

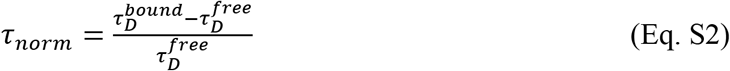

Both *τ_D_* and *τ_norm_* are listed in Table S5.

In contrast to other constructs, binding of the PRR to tubulin has a sigmoidal shape with a saturation plateau beyond 2 μM tubulin (Fig. S3). The binding curve fits to the Hill equation:

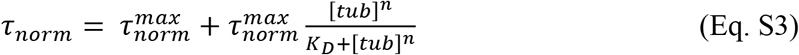

where 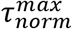 is the normalized diffusion time for tau:tubulin measured at 10 μM tubulin, *n* is the Hill coefficient and reflects the extent of cooperativity, *K_D_* is the apparent dissociation constant and [tub] is the concentration of tubulin. Fitting our data with this equation yields, *n* = 1.7 ± 0.2 and *K_D_* = 900 nM ± 10 nM. In our previous work, we used the engineered protein construct RB3, which binds tubulin with 1:2 RB3: tubulin dimer stoichiometry to determine the expected τ_D_ of a 1:2 protein:tubulin complex (3, 6). Here, the τ_D_ measured for the PRR at 10 μM tubulin (0.82 ± 0.03 ms) is consistent with a 1:2 tau:tubulin dimer stoichiometry. This observation, coupled with the cooperativity seen in Hill equation fit, strongly supports the presents of two tubulin-dimer binding sites in the PRR.

### FRET instrument and analysis

For FRET histograms where the protein signal was readily distinguishable from the ‘zero-peak’ (7), arising from imperfect labeling were carried out on our lab built instrument, as described above with a few modifications. The laser power is adjusted to ~30 μW (488nm diode-pumped solid-state laser, Spectra-Physics) prior to entering the microscope. Donor and acceptor photons were separated using a HQ585 long pass dichroic and further selected with ET525/50M band pass and HQ600 long pass filters (Chroma). For each path, the emission was focused onto the aperture of a 100 μm diameter optical fiber (OzOptics) coupled to an avalanche photodiode (Perkin-Elmer). Time traces were collected in 1 ms time bins for 1 hour. As described above, measurements were carried out in PEG-PLL coated Nunc chambers with 20-40 pM labeled tau following 5 minutes incubation with tubulin.

To differentiate photon bursts arising from transit of a labeled molecule from background fluorescence, a photon count threshold of 30 counts/ms was applied. For each burst, the energy transfer efficiency (ET_eff_) calculated using a lab-based written software (MATLAB) according to the following equation (8, 9):

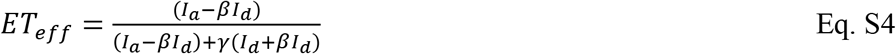

where *I_a_* is the intensity of the acceptor photons and *I_d_* is the intensity of the donor photons. Within our system, the bleed through of the donor channel into the acceptor channel (*β*) and the difference in the total quantum efficiency of the system and fluorophores (*γ*) were determined using Alexa Fluor 488 hydrazine (Invitrogen) and Alexa Fluor 594 hydrazine (Invitrogen) and fixed for analysis. Due to variation in instrument build and detector efficiency over the course of the study, *β* and *γ* were regularly re-determined and checked with DNA standards of 10, 14, and 18 bases labeled with Alexa Fluor 488 and Alexa Fluor 594 (Integrated DNA Technologies). The energy efficiencies were then binned, and the histograms fit using a sum of Gaussians in Origin. One Gaussian described the “zero-peak” (donor-only fluorescence) and the second peak described donor and acceptor labeled protein (main peak fit listed in Table S2). In some cases, the distribution was asymmetric (such as tau^17-149^). In these cases, three Gaussians were used to fit the data. The Gaussian that fit the dominant peak is reported in Table S2.

At some of the labeling positions, the proteins gave rise to low energy efficiencies with overlap with zero-peak, making it difficult to accurately determine the peak *ET_eff_* for the protein sample. To separate donor-only labeled species from the low energy, donor and acceptor labeled species, measurements were repeated on a commercial MicroTime 200 time-resolved confocal microscope (Picoquant) using its pulsed interleaved excitation FRET (PIE-FRET) mode. The power of the excitation lasers (485 nm and 562 nm) were matched ~30 μW at 40 MHz. The fluorescence emission was focused through a 100 μm pinhole and collected by avalanche photodiode. Fluorescence emission of the donor and acceptor fluorophores were separated using a HQ585 long pass dichroic and further selected with ET525/50M band pass and HQ600 long pass filters. SymphoTime 64 software was used to analyze the photon bursts to yield both the *ET_eff_* and stoichiometry factors for each burst, using photon threshold, binning and experimentally determined *β* and *γ* values as described previously. The binned histograms were fit as described above.

### Tubulin polymerization Assay

Polymerization of soluble tubulin was measured by monitoring the increase in scattered light at 340 nm. The tubulin was clarified as described above and buffer exchanged in phosphate buffer at pH 6.9 immediately prior to use. For polymerization reactions, 10 μM tubulin was incubated with 5 μM tau (for tau_2N_, tau_1N_, tau_0N_, and PRR-MTBR-R’) or 10 μM tau (for PRR, 2N-PRR, MTBR, and MTBR-R’) for 5 minutes on ice prior to the addition of 1 mM GTP. Immediately after the addition of GTP, the reaction was transferred to a warmed cuvette and the reaction was monitored for 10 minutes at 37 °C in a fluorometer (Fluorolog FL-1039/40, Horiba) with a photon counting module (SPEX DM302, Horiba) with both excitation and emission wavelengths set to 340 nm. Following polymerization, the samples were quickly returned to 4 °C; cold depolymerization is evidence that the proteins are not aggregated.

The curves were normalized to the brightest intensity within a given curve, and fit in Origin with:

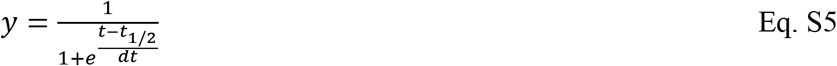

where y is the normalized fluorescence intensity, t is time, *t_1/2_* is the polymerization half-time and *dt* is the time constant. The mean and standard deviation of the *t_1/2_* values are listed in Table S4. The plotted graphs represent the average of the normalized triplicate with standard deviation.

---

**Fig. S1.**
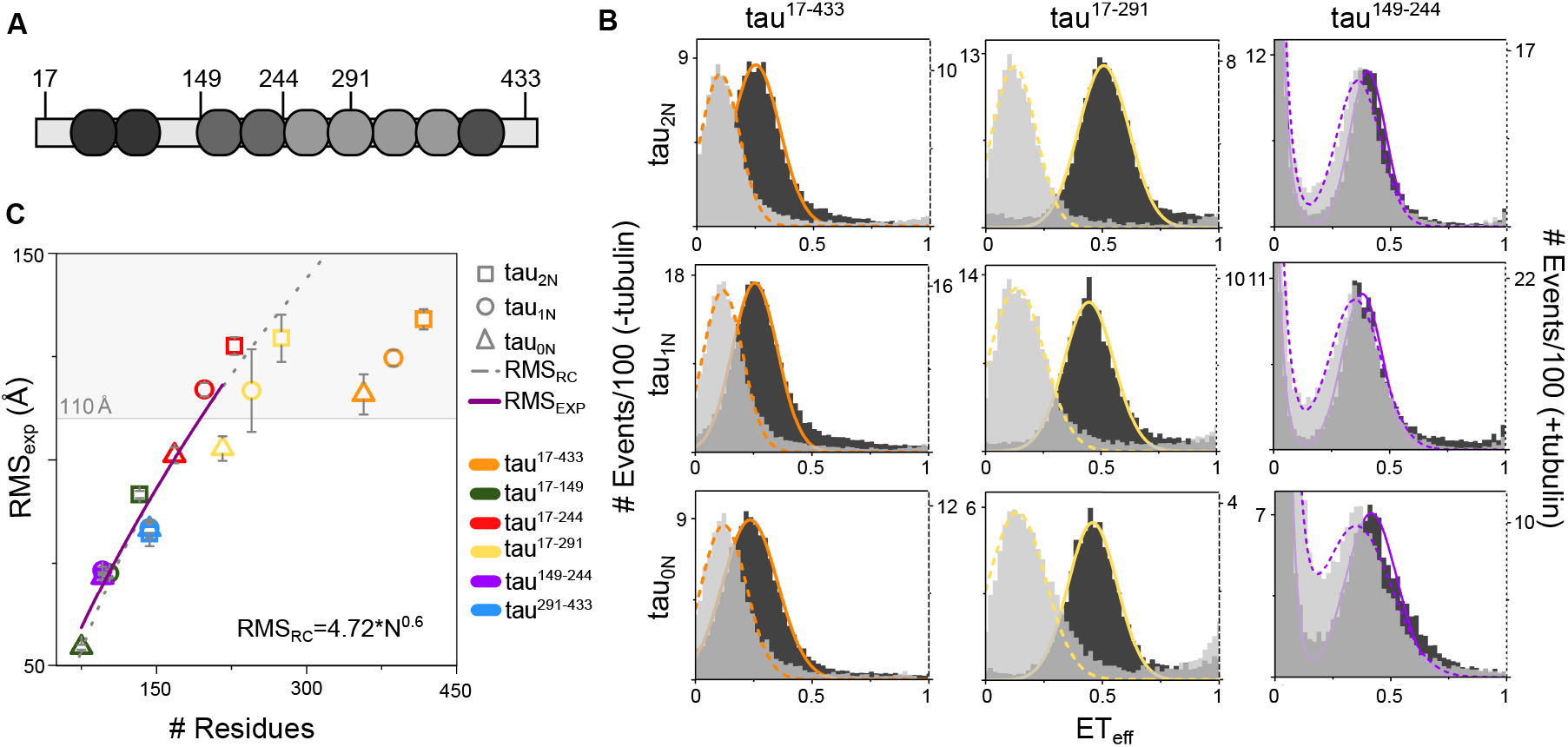
SmFRET of tau N-terminal isoforms. *(A)* Schematic of reference construct tau_2N_ with residues used for labeling for smFRET measurements are indicated. *(B)* SmFRET histograms of tau_2N_, tau_1N_ and tau_0N_ labeled at residues 17 and 433 (left), 17 and 291 (center), 149 and 244 (right) in the absence (dark: left axis) and presence (light: right axis) of 10 μM tubulin. Histograms for tau labeled at residues 291 and 433, 17 and 244, 149 and 244 are in Fig. 2. Fits to the data as a sum of Gaussian distributions as described in the *SI Appendix* are shown. *(C)* For measurements in the presence of tubulin, the root-mean-square distances (RMS) between labeling positions calculated from a Gaussian chain model (RMS_exp_) are plotted (10). Data are presented as mean ± SD, n≥3 independent measurements. For histograms with mean ET_eff_ values < 0.16 (shaded; corresponding to RMS_exp_ ≥ 110 Å), it is not possible to accurately calculate RMS_exp_ because the average distance between the fluorophores is to large for the dye pair used. For RMS_exp_ values < 110 Å, the data was fit using an exponential curve (RMS_exp_). For reference, the RMS calculated for a random coil (RMS_RC_) as in Reference (11) is indicated by the gray dashed line. See Table S2 for, numerical values of ET_eff_ mean ± SD, RMS_exp_ ± SD and RMS_RC_ for each construct.

**Fig. S2.**
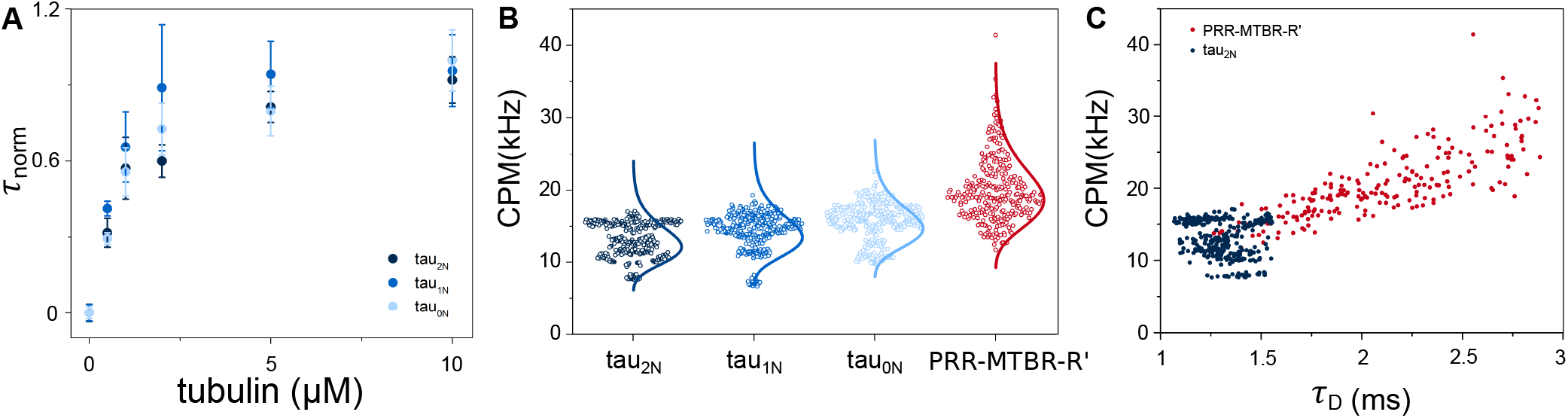
Relative binding and brightness of tau N-terminal isoforms. *(A)* The average τ_norm_ of tau constructs tau_2N_ (replotted from Fig. 2A for comparison), tau_1N_ and tau_0N_ are plotted against increasing tubulin concentration. Data are presented as mean ± SD, n≥3 independent measurements. See *SI Appendix* for details of data analysis. See Table S5 for numerical values for τ_D_ and τ_norm_ at 10 μM tubulin. *(B)* The corresponding CPM of individual autocorrelation curve fits of tau_2N_, tau_1N_, tau_0N_ and PRR-MTBR-R’ incubated with 10 μM tubulin from Fig. 3B with lognormal distribution overlaid. *(C)* Scatterplot of CPM versus τ_D_ of PRR-MTBR-R’ and tau_2N_ from panels *(B)* and Fig. 3B, respectively.

**Fig. S3.**
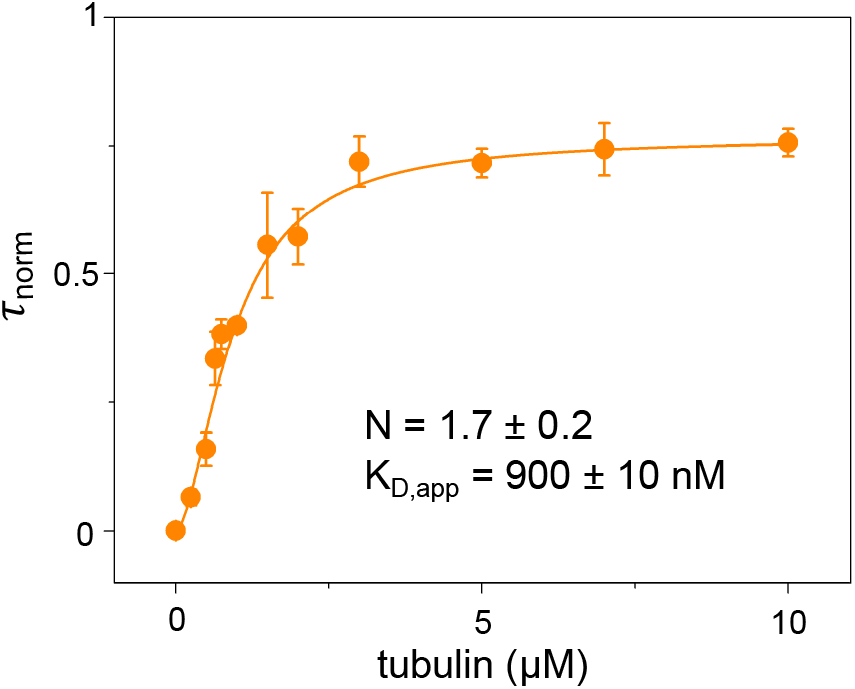
Cooperative binding by PRR. The average τ_norm_ and standard deviation of the PRR is plotted against increasing tubulin concentration (re-plotted from Fig. 4A with additional tubulin concentrations). The binding curve was fit to the Hill equation (Eq. S3) was to yield n=1.7 ± 0.2 and the apparent K_D_= 900 ± 10 nM. Data are presented as mean ± SD, n≥3 independent measurements. See *SI Appendix* for details of data analysis. See Table S5 for numerical values for τ_D_ and τ_norm_ at 10 μM tubulin.

**Fig. S4.**
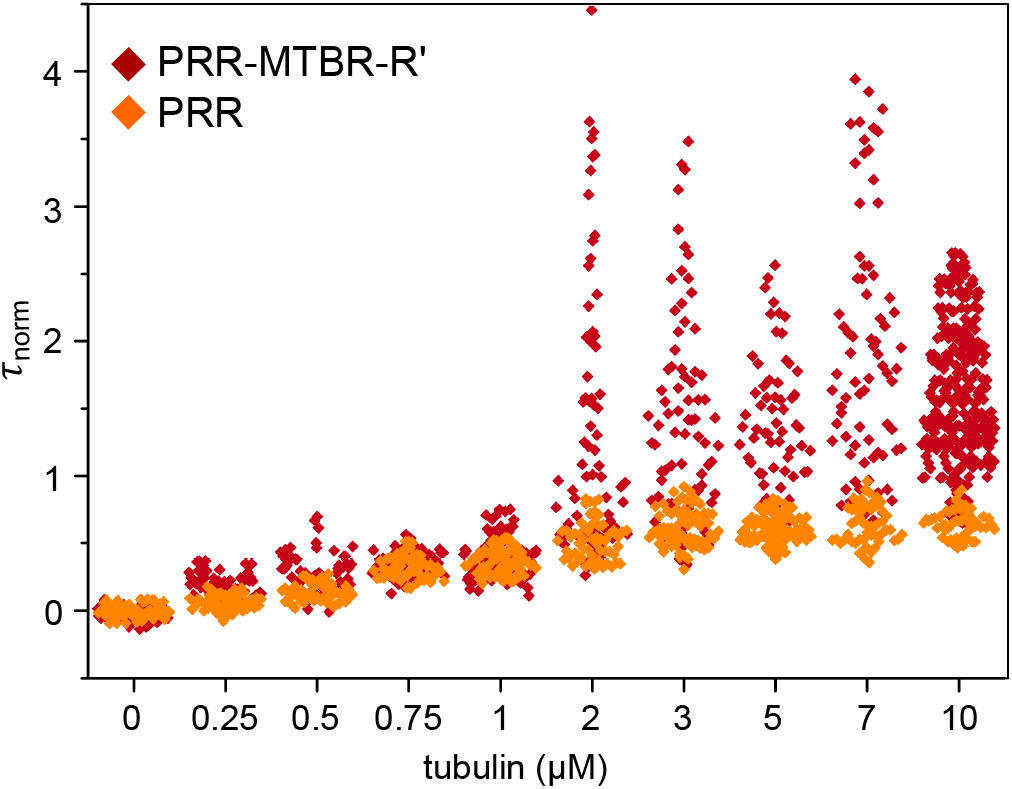
Comparison of heterogeneity of PRR and PRR-MTBR-R’. The *τ_D_* values from fitting individual autocorrelation curves for PRR-MTBR-R’ (from Fig. 3) and PRR (from Fig. 4) is plotted as a scatterplot against increasing tubulin concentration.

**Fig. S5.**
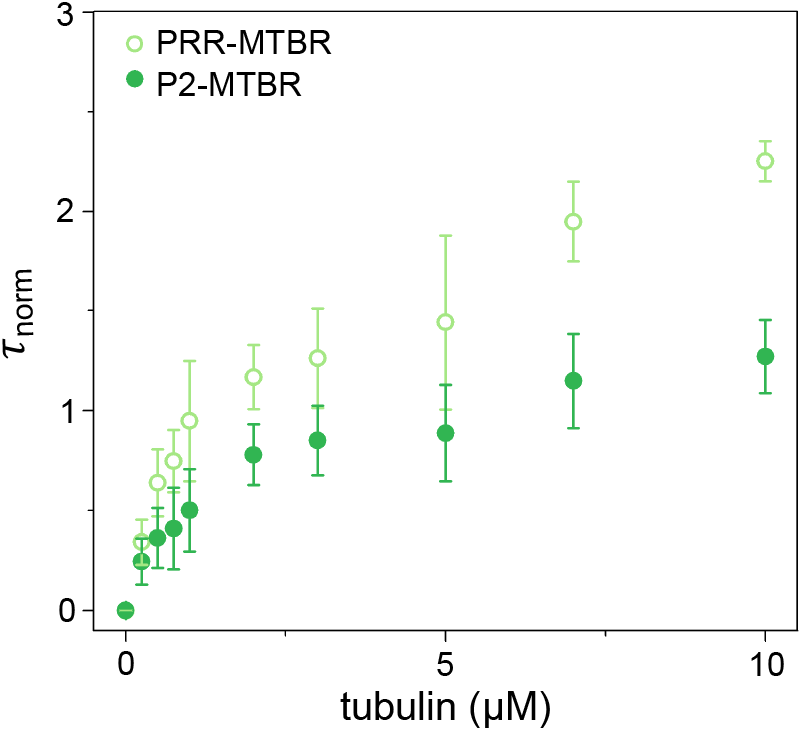
Impact of P1 on binding of the PRR-MTBR. The τ_norm_ of tau constructs PRR-MTBR and P2-MTBR measured by FCS are plotted against increasing tubulin concentration. Data are presented as mean ± SD, n≥3 independent measurements. See *SI Appendix* for details of data analysis. See Table S5 for numerical values for τ_D_ and τ_norm_ at 10 μM tubulin.

**Fig. S6.**
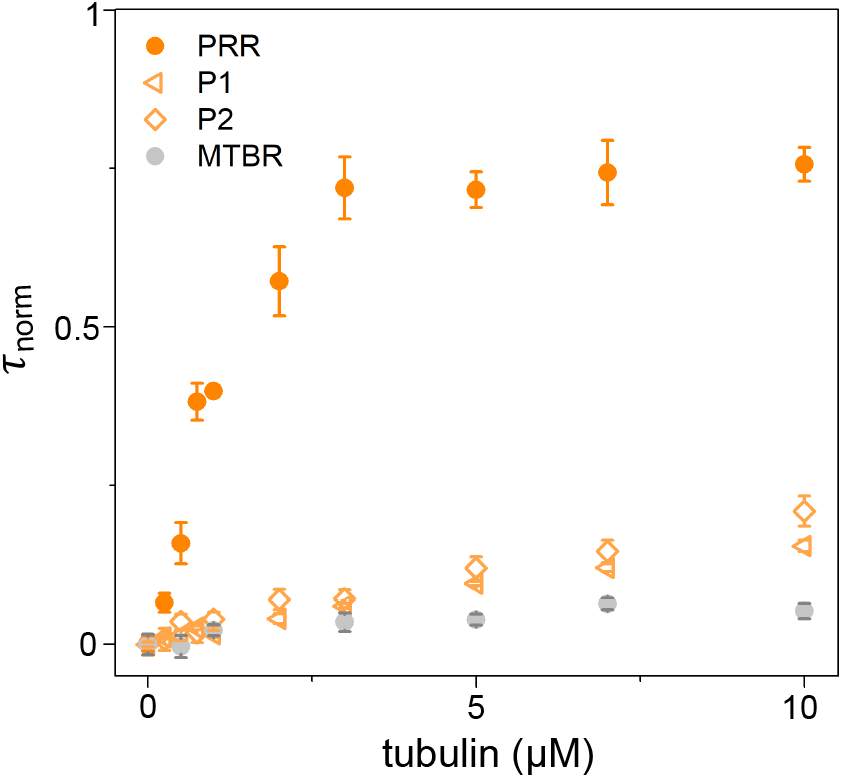
Relative binding of P1 and P2. The average τ_norm_ and standard deviation of FCS tau constructs P1 and P2 are plotted against increasing tubulin concentration. Data are presented as mean ± SD, n≥3 independent measurements. Binding curves of the PRR and MTBR (both from Fig. 4A) are plotted for comparison. See *SI Appendix* for details of data analysis. See Table S5 for numerical values for τ_D_ and τ_norm_ at 10 μM tubulin.

**Fig. S7.**
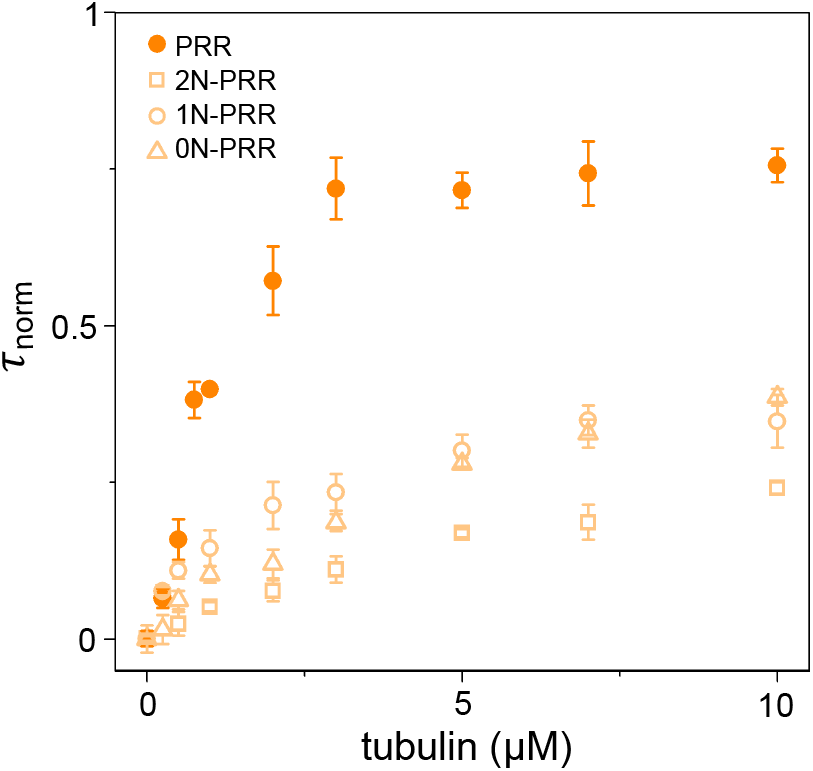
Impact of N-terminal inserts on the binding of PRR. The τ_norm_ of tau constructs 1N-PRR and 0N-PRR are plotted against increasing tubulin concentration. Tau constructs PRR and 2N-PRR are re-plotted from Fig. 4A for comparison. Data are presented as mean ± SD, n≥3 independent measurements. See *SI Appendix* for details of data analysis. See Table S5 for numerical values for τ_D_ and τ_norm_ at 10 μM tubulin.

**Fig. S8.**
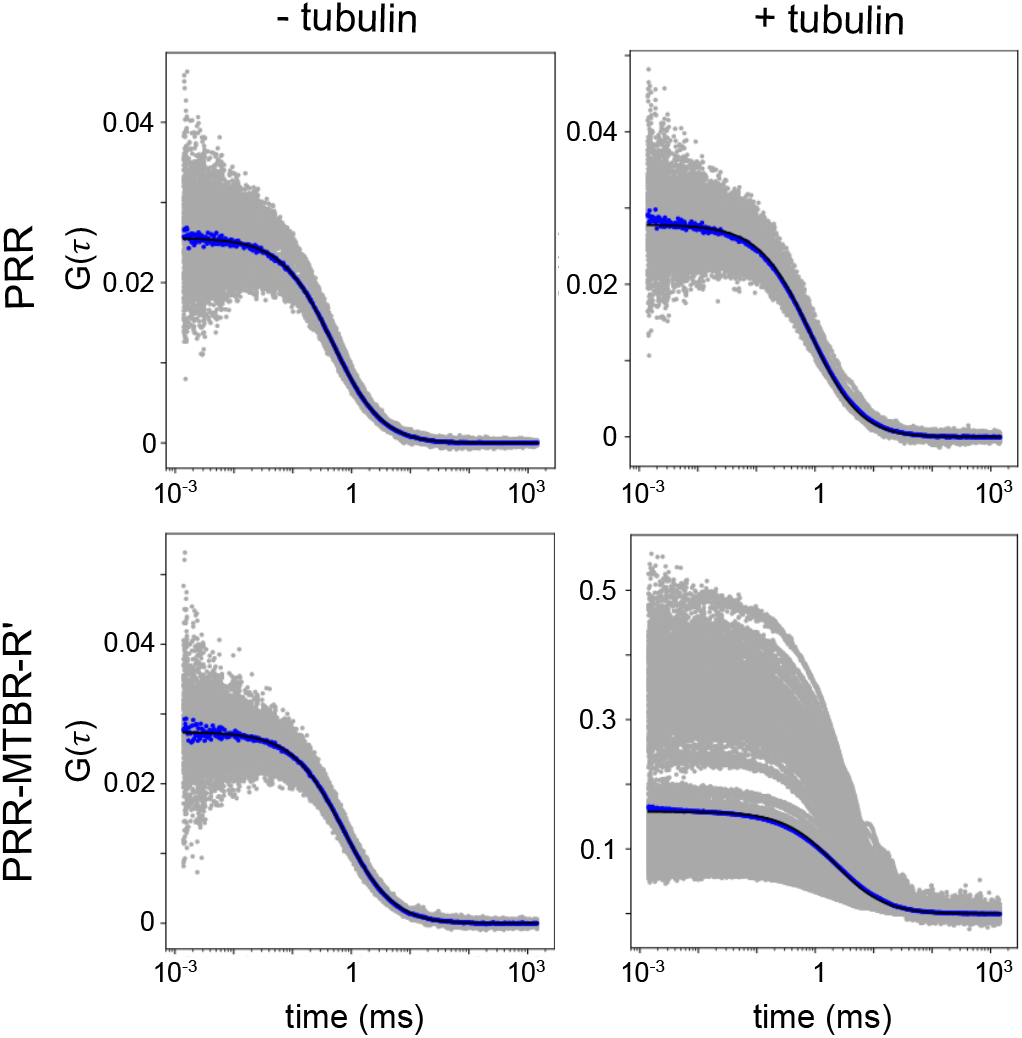
Heterogeneity in tubulin-bound PRR-MTBR-R’. Individual autocorrelation curves (gray dots) are plotted for PRR (upper) and PRR-MTBR-R’ (lower) in the absence (left) or presence (right) of 10 μM tubulin. Averaged curves are shown with blue dots and fits of the averaged curves to Eq. S1 are in black.

**Fig. S9.**
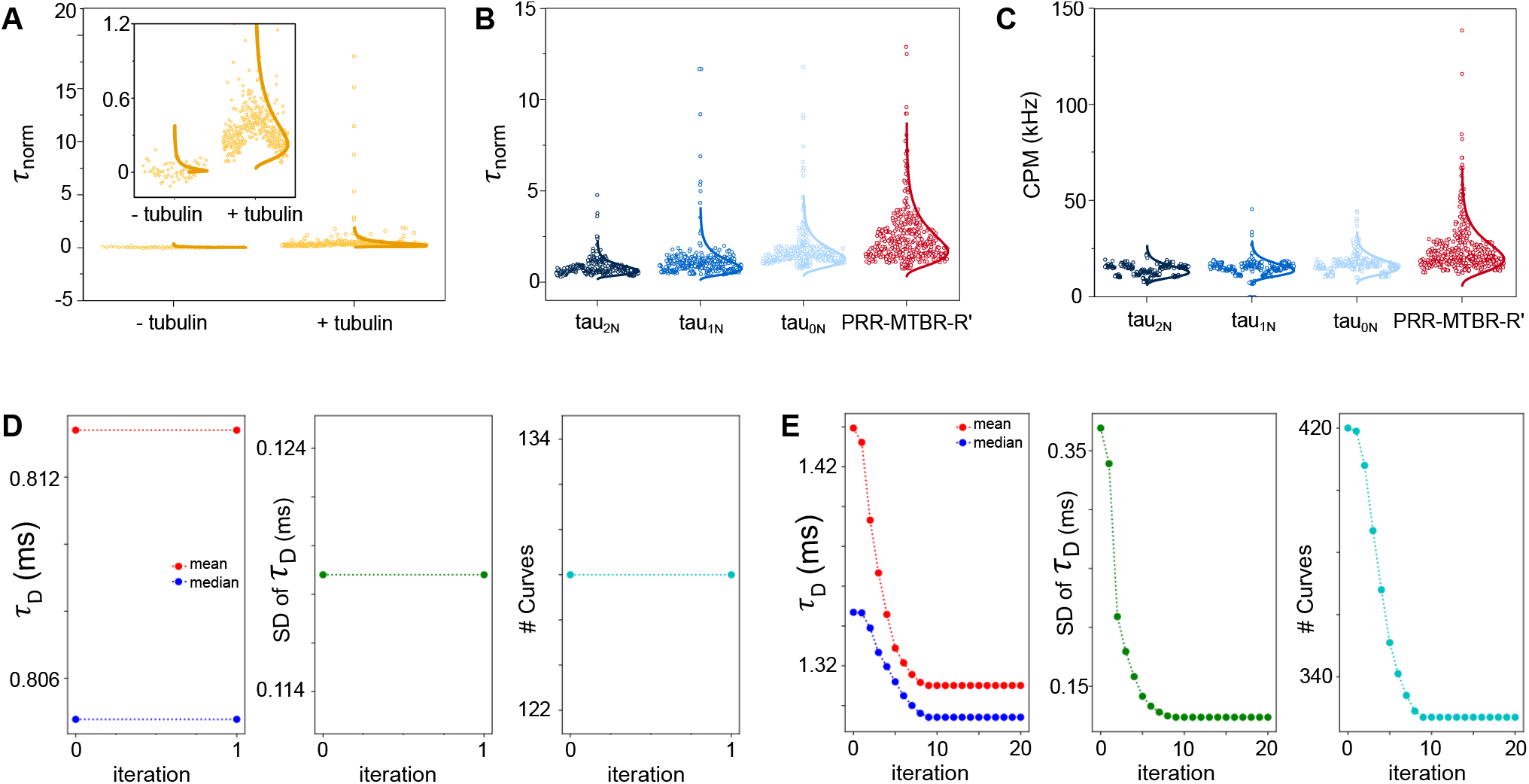
Filtering algorithm. *(A)* τ_norm_ of MTBR-R’ in the absence or presence of tubulin without any filtering. Inset is a magnification of the y-axis for low τ_norm_. *(B)* τ_norm_ of tau_2N_, tau_1N_, tau_0N_ and PRR-MTBR-R’ with 10 μM tubulin Fig. 4A after goodness-of-fit filtering, but without iterative filtering (as described in the *SI Appendix*) with lognormal distribution overlaid. *(C)* The corresponding CPM (kHz) for each point in *(B) (D)* Mean and median τ_D_ (right), the standard deviation of τ_D_ (middle) and number of curves (left) for tau_2N_ over the iteration number (1) for tau_2N_ in the absence of tubulin. No curves are selected for elimination in this tau-only sample. *(E)* Mean and median τ_D_ (right), the standard deviation of τ_D_ (middle) and number of curves (left) for tau_2N_ in the presence of 10 μM tubulin over the iteration number demonstrating convergence of the algorithm (at iteration 9) described in the *SI Appendix*.

**Table S1.** Tau constructs and their corresponding residues.

| **construct** | **residues** | **C<sub>FCS</sub>** |
| --- | --- | --- |
| tau<sub>2N</sub> | 1-441 | 433 |
| tau<sub>1N</sub> | 1-441<sub>Δ45-75</sub> | 433 |
| tau<sub>0N</sub> | 1-441<sub>Δ45-102</sub> | 433 |
| tau<sub>ΔN</sub> | 148-441 | 433 |
| tau<sub>ΔC</sub> | 1-395 | 17 |
| 2N | 1-150 | 17 |
| PRR | 148-244 | 149 |
| MTBR | 244-372 | 322 |
| MTBR-R' | 244-395 | 244 |
| PRR-MTBR-R' | 148-395 | 149 |
| PRR-MTBR | 148-372 | 149 |
| 2N-PRR | 1-244 | 17 |
| 1N-PRR | 1-244<sub>Δ45-75</sub> | 17 |
| 0N-PRR | 1-244<sub>Δ45-102</sub> | 17 |
| 2N-MTBR-R' | 1-395<sub>Δ151-253</sub> | 17 |
| P1 | 148-197 | 149 |
| P2 | 198-244 | 244 |
| P2-MTBR | 198-372 | 322 |

All numbering throughout the manuscript is based on tau<sub>2N</sub>. Unless otherwise noted, all constructs contain C291S and C322S mutations. C<sub>FCS</sub> is the residue number mutated to cysteine for labeling for FCS measurements.

**Table S2.** SmFRET of N-terminal isoforms

| isoform | labels | # residues | ET<sub>eff</sub> |  | RMS<sub>exp</sub> (Å) |  | RMS<sub>RC</sub> (Å) |
| --- | --- | --- | --- | --- | --- | --- | --- |
|  |  |  | - tubulin | + tubulin | - tubulin | + tubulin |  |
| **tau<sub>2N</sub>** | 17 433 | 417 | 0.27 ± 0.02 | 0.10 ± 0.01 | 88 ± 3 | 134 ± 2 | 181 |
|  | 17 291 | 275 | 0.52 ± 0.02 | 0.11 ± 0.02 | 62 ± 1 | 129 ± 6 | 141 |
|  | 17 244 | 228 | 0.62 ± 0.01 | 0.11 ± 0.01 | 52 ± 1 | 127 ± 2 | 126 |
|  | 17 149 | 133 | 0.48 ± 0.01 | 0.25 ± 0.01 | 64 ± 1 | 92 ± 1 | 91 |
|  | 149 244 | 96 | 0.43 ± 0.01 | 0.39 ± 0.01 | 68 ± 1 | 72 ± 1 | 75 |
|  | 291 433 | 143 | 0.44 ± 0.01 | 0.31 ± 0.02 | 67 ± 1 | 82 ± 3 | 95 |
| **tau<sub>1N</sub>** | 17 433 | 386 | 0.26 ± 0.01 | 0.12 ± 0.01 | 89 ± 1 | 125 ± 2 | 172 |
|  | 17 291 | 245 | 0.45 ± 0.01 | 0.14 ± 0.02 | 66 ± 2 | 117 ± 10 | 131 |
|  | 17 244 | 198 | 0.58 ± 0.01 | 0.14 ± 0.01 | 56 ± 1 | 117 ± 2 | 115 |
|  | 17 149 | 103 | 0.56 ± 0.01 | 0.39 ± 0.01 | 57 ± 1 | 73 ± 1 | 78 |
|  | 149 244 | 96 | 0.41 ± 0.01 | 0.38 ± 0.01 | 70 ± 1 | 73 ± 1 | 75 |
|  | 291 433 | 143 | 0.46 ± 0.01 | 0.30 ± 0.01 | 65 ± 1 | 83 ± 1 | 95 |
| **tau<sub>0N</sub>** | 17 433 | 358 | 0.28 ± 0.02 | 0.15 ± 0.01 | 86 ± 1 | 115 ± 5 | 165 |
|  | 17 291 | 216 | 0.46 ± 0.01 | 0.19 ± 0.01 | 67 ± 1 | 103 ± 3 | 122 |
|  | 17 244 | 168 | 0.61 ± 0.01 | 0.20 ± 0.01 | 54 ± 1 | 101 ± 2 | 105 |
|  | 17 149 | 75 | 0.72 ± 0.01 | 0.60 ± 0.01 | 46 ± 1 | 54 ± 1 | 65 |
|  | 149 244 | 96 | 0.44 ± 0.01 | 0.40 ± 0.01 | 68 ± 1 | 71 ± 1 | 75 |
|  | 291 433 | 143 | 0.46 ± 0.01 | 0.30 ± 0.02 | 65 ± 1 | 83 ± 2 | 95 |

**Table S3.** Descriptive statistics of tau:tubulin

| construct | pre-filtering |  |  |  | post-filtering |  |  |  |
| --- | --- | --- | --- | --- | --- | --- | --- | --- |
| | median | $\tau_D$ (ms)<br>mean ± SD | IQR | # curves | median | $\tau_D$ (ms)<br>mean ± SD | IQR | # curves |
| PRR-MTBR-R' | 2.24 | 2.52 ± 1.41 | 0.99 | 394 | 2.02 | 2.06 ± 0.42 | 0.63 | 304 |
| tau<sub>2N</sub> | 1.35 | 1.43 ± 0.34 | 0.29 | 419 | 1.29 | 1.31 ± 0.12 | 0.23 | 327 |
| tau<sub>1N</sub> | 1.55 | 1.73 ± 1.31 | 0.40 | 424 | 1.50 | 1.51 ± 0.20 | 0.34 | 383 |
| tau<sub>0N</sub> | 1.61 | 1.80 ± 0.86 | 0.47 | 434 | 1.55 | 1.57 ± 0.25 | 0.36 | 378 |

Statistics of the diffusion times from select tau constructs incubated with 10 μM tubulin without and with our filtering algorithm. IQR and SD stand for interquartile range and standard deviation respectively. See SI Appendix for details.

**Table S4.** Tau-mediated Polymerization

| **construct** | **$t_{1/2}$ (s)** | **tau : tubulin** |
| --- | --- | --- |
| PRR-MTBR-R' | $52 \pm 7$ | 1:2 |
| tau<sub>2N</sub> | $137 \pm 9$ | 1:2 |
| tau<sub>1N</sub> | $88 \pm 13$ | 1:2 |
| tau<sub>0N</sub> | $76 \pm 10$ | 1:2 |
| PRR | $96 \pm 55$ | 1:1 |

**Table S5.**
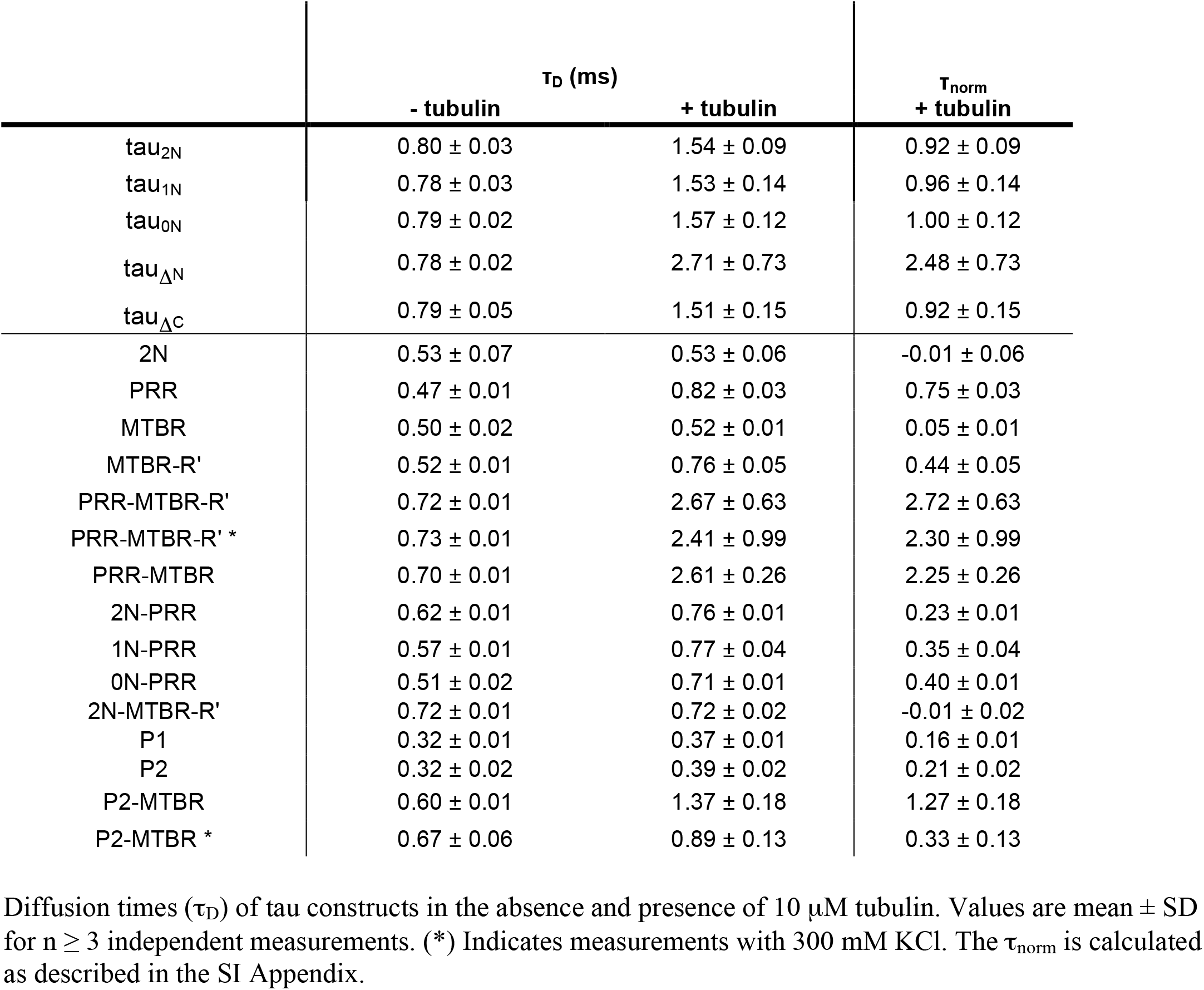
Summary of FCS diffusion times

**Table S6.**
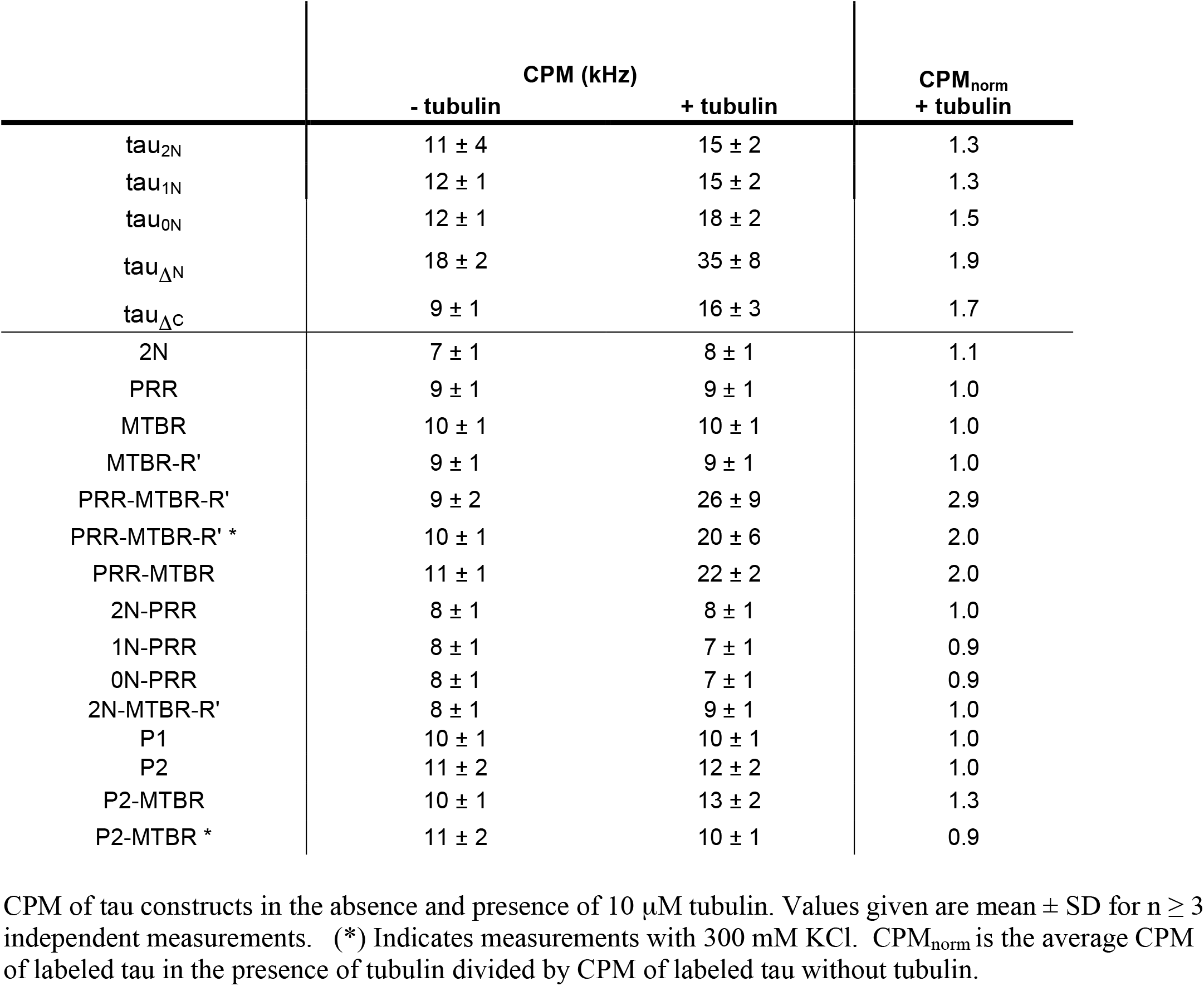
Summary of CPM from FCS measurements

|  | CPM (kHz) |  | CPM<sub>norm</sub><br>+ tubulin |
| --- | --- | --- | --- |
|  | - tubulin | + tubulin |  |
| tau<sub>2N</sub> | 11 ± 4 | 15 ± 2 | 1.3 |
| tau<sub>1N</sub> | 12 ± 1 | 15 ± 2 | 1.3 |
| tau<sub>0N</sub> | 12 ± 1 | 18 ± 2 | 1.5 |
| tau<sub>ΔN</sub> | 18 ± 2 | 35 ± 8 | 1.9 |
| tau<sub>ΔC</sub> | 9 ± 1 | 16 ± 3 | 1.7 |
| 2N | 7 ± 1 | 8 ± 1 | 1.1 |
| PRR | 9 ± 1 | 9 ± 1 | 1.0 |
| MTBR | 10 ± 1 | 10 ± 1 | 1.0 |
| MTBR-R' | 9 ± 1 | 9 ± 1 | 1.0 |
| PRR-MTBR-R' | 9 ± 2 | 26 ± 9 | 2.9 |
| PRR-MTBR-R' * | 10 ± 1 | 20 ± 6 | 2.0 |
| PRR-MTBR | 11 ± 1 | 22 ± 2 | 2.0 |
| 2N-PRR | 8 ± 1 | 8 ± 1 | 1.0 |
| 1N-PRR | 8 ± 1 | 7 ± 1 | 0.9 |
| 0N-PRR | 8 ± 1 | 7 ± 1 | 0.9 |
| 2N-MTBR-R' | 8 ± 1 | 9 ± 1 | 1.0 |
| P1 | 10 ± 1 | 10 ± 1 | 1.0 |
| P2 | 11 ± 2 | 12 ± 2 | 1.0 |
| P2-MTBR | 10 ± 1 | 13 ± 2 | 1.3 |
| P2-MTBR * | 11 ± 2 | 10 ± 1 | 0.9 |

CPM of tau constructs in the absence and presence of 10 μM tubulin. Values given are mean ± SD for $n \geq 3$ independent measurements. (\*) Indicates measurements with 300 mM KCl. CPM<sub>norm</sub> is the average CPM of labeled tau in the presence of tubulin divided by CPM of labeled tau without tubulin.

